# “Genetic structure and sodium channel gene mutation analysis related to pyrethroid insecticide toxicity in sylvatic Andean *Triatoma infestans* from Bolivia”

**DOI:** 10.1101/512392

**Authors:** Paula L. Marcet, Pablo Santo-Orihuela, Louisa A. Messenger, Claudia V. Vassena

## Abstract

**BACKGROUND:** Sylvatic populations of *Triatoma infestans* represent a challenge for vector control as these populations are not targeted by control activities and could play a key role in post-spraying house re-infestation. Improved understanding of sylvatic foci, population distribution, dispersion patterns, gene flow between sylvatic and domestic populations, as well as characterization of insecticide resistance profiles, is crucial to optimize vector control interventions.

**METHODS:** We analyzed the genetic relationship of five Andean populations from Bolivia from localities with distinct insecticide susceptibility profiles (sylvatic: 20 de Octubre, Illicuni, Kirus Mayu and Mataral and one domestic from Mataral). Individual multilocus genotypes based on 8 microsatellites and the DNA sequence of a fragment of the cytochrome B (cytB) gene were obtained for 92 individuals. We compared the cytB haplotypes with previously reported Andean *T. infestans* haplotypes and evaluated the directionality and possibly history of gene flow among populations. Each specimen was screened for 2 nucleotide mutations (L1014 and L9251) of the sodium channel gene (*kdr*), described for *T. infestans* and related to pyrethroid resistance.

**RESULTS:** Significant genetic differentiation was observed among all populations, reflecting current genetic isolation among them. However, individuals of admixed origin were detected in four populations, especially between the sylvatic and domestic populations from Mataral. Historical analysis of gene flow suggests that insecticide resistance is conferred by ancient trait(s) in *T. infestans* sylvatic populations that are capable of invading domiciles. The *kdr* mutation L1014 was identified in one individual from Mataral, while the L9251 mutation was not detected in any population. The low frequency of *kdr* mutations in these populations suggests this mechanism is unlikely to be the primary cause of the observed altered insecticide susceptibility. However, the resistance conferring mutation is present in the area and with the potential to be selected under insecticidal pressure.

**CONCLUSIONS:** These results emphasize the need for stronger entomological surveillance in the region, including early detection of house invasion, particularly post-spraying, monitoring for resistance to pyrethroids and the design of integrative control actions that consider both sylvatic foci around domestic settings as well as the bug dispersion dynamics.

## INTRODUCTION

Chagas disease is caused by *Trypanosoma cruzi*, a protozoan parasite that infects humans as well as domestic and sylvatic mammals. This disease can be transmitted by approximately 150 species of hemipteran insect vectors of which *Triatoma infestans* (Klug 1834) (Hemiptera: Reduviidae, Triatominae) is the most important in the southern countries of Latin America. [1, 2]. The most recent reports estimate that Chagas disease affects approximately 6 million people worldwide; approximately 70 million people are at risk of infection in 21 Latin American countries [3].

Domiciliary populations of *T. infestans* have been successfully controlled by spraying with pyrethroid insecticides in parts of southern South America [1, 4]; this contributed to the reduction in the geographical distribution of *T. infestans* from an estimated 6.28 million km^2^ in the 1960s to estimations of less than 1 million km2 today [1, 5]. Moreover, several countries and regions in South America, have been certified as having interrupted domestic vector transmission [6].

The “Gran Chaco Region”, which covers 1.3 million km^2^ including territories in Bolivia, Paraguay and Argentina, is highly endemic for Chagas disease transmission; it has proven to be particularly difficult to control domestic infestation by *T. infestans* in the region. Rapid house reinfestation post-spraying and insecticide resistance are among the factors associated with vector control failure in parts of this region [7–11].

*T. infestans* has been found in sylvatic foci in several areas in the Chaco region, some reportedly invading houses and are associated with *T. cruzi* transmission to humans [12–23]. Sylvatic populations are not targeted by control interventions and could re-invade houses treated with insecticide, especially if the population has resistant variants that can be selected under insecticidal pressure. Remarkably, certain sylvatic populations from Bolivia have shown low susceptibility to pyrethroid insecticides [24], and some also exhibited a lower sensitivity to fenitrothion [25].

Pyrethroid resistance in *T. infestans* was initially described in Argentina and Bolivia in the late 1990s. It later became implicated in failure of local control programs, where certain populations from the Gran Chaco area required up to 1000 times the amount of insecticide to kill the same proportion of susceptible individuals [26]. Increasing reports indicate that several natural populations are resistant to insecticides, and significant variability in susceptibility levels has been observed among them [27–30].

Resistance traits are considered pre-adaptive, implying that there are a few resistant insects in each natural population, which would survive under the selective pressure of insecticide exposure, selecting for the resistant genetic variants that would be inherited and in turn, become more frequent in subsequent generations. The dispersal rate of resistant traits will depend on pre-existing genetic variation in natural populations, selection intensity, fitness cost associated to expressing the resistant variant, local adaptation and levels of gene flow among populations. Such resistance dynamics have been reported in populations of other medically-important vector species, particularly *Aedes aegypti* [31, 32]. Following this rationale, three general assumptions can be made regarding population genetic diversity and insecticide resistance:

a. Insecticide resistant triatomine populations are expected to be less genetically diverse than a wild-type population as they have likely undergone a period of selective pressure (i.e. insecticide exposure).
b. Sylvatic populations are expected to be more genetically diverse than domestic populations as they are in theory, ancient progenitors, that have accumulated diversification over time, in the absence of selective pressure, as they are not being targeted by vector control activities.
c. Domestic populations are expected to have been the subject of greater selection pressures, compared to their sylvatic counterparts and therefore be much less diverse.

Elucidating population genetic dynamics through high resolution genetic markers in the context of well characterized insecticide resistance profiles could test these hypothesis and provide invaluable information for an integrated vector control approach. Establishing the role of sylvatic populations of *T. infestans* as potential sources of re-infestation and understanding the impact of insecticide resistance on triatomine population genetics is crucial to safeguard the continued effectiveness of current control efforts. Previous studies using high resolution molecular markers demonstrated that genetic differentiation is detectable up to the capture site level (local populations) if sufficient numbers of bugs are captured per site to constitute a representative population for genetic analyses comparisons [33–36].

The aim of this work was to establish and characterize the genetic profiles of four sylvatic and one domestic Andean populations of *Triatoma infestans* from Bolivia, and to interpret their genetic relationships in the context of their differing levels of insecticide susceptibility. Moreover, an evaluation of the mechanisms likely responsible for the pyrethroid resistance observed in these populations including esterase enzymes and Ыr-mutations is discussed.

## MATERIALS AND METHODS

### Insect origin

Collection methods were described in detail in [24]. Briefly, *T. infestans* populations were sampled from domiciliary areas in Mataral, Department of Cochabamba, Bolivia (hereby named Mataral-D) and a sylvatic area located approximately 2 km away (Mataral-S). Sylvatic bugs were also collected from sites in Cochabamba (Ilicuni and 20 de Octubre) and Potosí (Kirus Mayu). Sylvatic triatomines were sampled from rock-piles using mouse-baited sticky traps [37]. From each population a colony was established with a minimum of 10 founder individuals, and maintained at 28±1ºC, 50% RH, and a photoperiod of 12:12 (L:D) h at the Centro de Investigaciones de Plagas e Insecticidas (CIPEIN, CITEFA-CONICET), Buenos Aires, Argentina. Rearing conditions are described in detail elsewhere [38, 39]. For this study, bugs from each population from the second or third colony generations were used. Genomic DNA was obtained from three legs of each bug using the Wizard Genomic Purification Kit (Promega^®^) following the manufacturer recommendations.

The insecticide resistance profile of these populations was established by toxicological bioassays performed according to World Health Organization [40] standard protocols [24].

### Multilocus microsatellite genotyping

A multilocus genotype was obtained for each individual based on nine microsatellite loci, developed and optimized for *T. infestans* following the amplification conditions previously described [33, 41]. Multiplex PCR reactions were carried out with 2 pairs of loci with the same annealing temperature and different dye colors (Tinf_ms5 with Tinf_ms42 and Tinf_ms22 with Tinf_ms27). DNA fragment detection with 1bp resolution was performed with an automated DNA sequencer (ABI 3130, Applied Biosystems) and size determination was obtained with GeneMapper 4.1 (Applied Biosystems). Ninety-two colony-reared, second-third generation individuals were sampled for microsatellite genotyping (Table 1).

**Table 1:** Number of bugs per population evaluated per molecular marker: msat= microsatellites; cytB= cytochrome B; **kdr**: evaluated for 2 mutations in the sodium channel gene (L1014 and L9251). Toxicity of topically applied deltamethrin in first instars nymphs of *T. infestans* from Bolivia (^1^Data previously published [24]). *Susceptible strain reared at CIPEIN. **LD50**: Lethal dose to kill 50% of the sample measured in ng/insect; **CL**= confidence limit; **RR**: resistance ratio.

| Collection site | msat | cytB | kdr | LD50<sup>1</sup><br>(95% CL) | RR<sup>1</sup><br>(95% CL) |
| --- | --- | --- | --- | --- | --- |
| NFS* | - | - | - | 0.13<br>(0.11-0.15) | - |
| Mataral-D<br>(Mat-D) | 19 | 19 | 13 | 2.25<br>(0.28-4.80) | 17.4<br>(11.88-25.43) |
| Mataral-S<br>(Mat-S) | 11 | 11 | 6 | 1.53<br>(0.53-3.34) | 11.9<br>(9.43-14.93) |
| Kirus Mayu<br>(KM) | 22 | 23 | 15 | 0.95<br>(0.49-1.66) | 7.4<br>(5.78-9.25) |
| Ilicuni<br>(Ili) | 18 | 18 | 16 | 0.25<br>(0.14-0.39) | 1.9<br>(1.44-2.56) |
| 20 de Octubre<br>(20oct) | 22 | 22 | 17 | 0.88<br>(0.08-1.84) | 6.8<br>(4.98-9.26) |

### Microsatellite analysis

Allele number per locus and population were obtained directly after binning, and mean allele number among all loci were compared. The inbreeding coefficient **Fis** [42]; allelic richness **Ar** [43], gene diversity or expected heterozygosity (He) and observed heterozygosity (Ho) [44] were obtained with the Excel add-in MS_tools.xla for Microsoft Excel or with *F*_ST_AT2.9.3.2 [45]. Bonferroni correction was applied to determine p-values for multiple comparisons [46]. The fit to Hardy-Weinberg (HW) equilibrium expectations was evaluated using the U score test available in Genepop 4.2, under the null hypothesis of random union of gametes. Sample-size-corrected private (population-specific) allele frequency per locus (PA/L) was calculated in HP-RARE [47]. Multilocus linkage disequilibrium, estimated by the index of association (*I*_A_), was calculated in MULTILOCUS 1. 3b [48] and statistical significance was evaluated by comparison with a null distribution of 1000 randomizations.

Pairwise *F*_ST_ values and significance levels were calculated with ARLEQUIN 3.01 as in Weir & Cockerham [42]. Fisher exact test for population differentiation comparing genic and genotypic frequencies was implemented in Genepop 4.2. Population clustering was explored using a neighbor-joining (NJ) tree based on pairwise distances (D_AS_: 1 proportion of shared alleles at all loci/n). A Mantel’s test for the effect of isolation by distance within populations (pairwise genetic vs. geographic distance) was performed in GenAlEx 6.5 using 10,000 random permutations [49]. A Hierarchical Analysis of Molecular Variance (AMOVA) [50] was performed at three levels of population structure: among communities, among populations and among individuals in a population, in ARLEQUIN 3.01 [51].

## Population structure

Population genetic structure was examined using the Bayesian model-based approach [52–54] implemented in STRUCTURE 2.3.4. Each individual MLG in the sample is probabilistically assigned to one of K populations, or jointly to two or more populations if their genotypes might have an admixed origin. Simultaneously, the method determines the number of significant K genetic clusters within the total sample. The number of clusters evaluated (=K) ranged from 1 to 10. The analysis was performed using 35 replicate runs per K value, a burn-in period length of 50,000 and a run length of 50,000. The analysis model used was the admixed and correlated allele frequencies, with no prior information on the origin of the individuals. The final selection of the sample K value was based on the log-probability of the data between successive K values [55] implemented in the online version of STRUCTURE HARVESTER [56].

## Mitochondrial (mtDNA) analysis

The cytB gene fragment was amplified as previously described [57] for a total of 74 specimens from the five populations (Table 1). SeqManPro Software 14.1 (DNAstar) was used to assemble and align forward (F) and reverse (R) chromatograms, and obtain the consensus sequence for each individual. MEGA 5 software [58] was used to align multiple sequences and to examine the phylogenetic relationships among sequences. Standard genetic variability (haplotype diversity, Hd and nucleotide diversity: π) and differentiation among sequences were evaluated using DnaSP version 5.1 software [59]. A Nexus matrix was constructed for haplotype network analysis in PopART using a median-joining model based on 1000 iterations with default parameters [60, 61].

To evaluate the possible directionality and history of gene flow among populations, we compared the haplotypes obtained in this work with previously reported Andean *T. infestans* haplotypes. Sequences were trimmed from 666bp to 388bp and a second haplotype network was built using the same software and criteria including 25 sequences from sylvatic Bolivian *T. infestans* previously deposited in GenBank: HQ333215-HQ333239 [15, 62].

### Screening for mutations in the *kdr* gene

A fragment of the DNA sequence of the sodium channel gene was amplified and screened for two nucleotide mutations previously reported in *T. infestans* associated with pyrethroid knockdown resistance (*kdr*) [63, 64]. The protocol has been previously described [65] and was used to screen 67 individuals from the five populations (Table 1).

## Results

### Population diversity analysis

Ninety-two individual *T. infestans* samples were grouped *a priori* into five populations based on their geographical origin (Table 1): Mataral domestic (Mat-D), Mataral sylvatic (Mat-S), Ilicuni (Ili), 20 de Octubre (20oct) and Kirus Mayu (KM) and were genotyped across eight polymorphic microsatellites. In contrast with genotyping data from *T. infestans* in Argentina and Paraguay, the locus Tims_64 did not amplify consistently across samples. In addition, locus Tims_65 did not amplify in either domestic or sylvatic Mataral populations, indicative of null alleles; therefore, neither loci were utilized in the final population analysis.

A total of 84 unique individual multilocus genotypes (MLG) were identified across populations (Table 2). Genetic diversity levels were high and consistent across most populations as evidenced by similar frequencies of unique MLGs, levels of allelic richness (3.1-5.4); distance between shared alleles (0.38-0.67); private alleles per locus (0.42-1.26) and average gene diversity (0.44-0.74) (Table 2). Only minor differences were observed between populations with larger numbers of founding individuals (e.g. 20oct and KM), indicating minimal genetic diversity was lost by culturing population members prior to genetic analysis.

Although it had the lowest number of individuals available for testing, the sylvatic population from Mataral displayed the highest parameters related to genetic diversity (average number of alleles = 5.5, Ar =5.37, He = 0.74, av PIC = 0.65). Moreover, individuals from this area demonstrated the highest number of private alleles per locus (1.26) and Fis value (0.22) indicative that this may be an ancestral population with restricted intra-population gene flow (i.e. sub-structure), respectively (Table 2). The populations 20oct, KM and Mat-S presented significant deviations from HW equilibrium, and all due to significant heterozygote deficits.

**Table 2.** Population genetic parameters for Bolivian T. infestans based on eight microsatellite loci. **#MGL8/N**: number of complete multilocus genotype for 8 loci / number of individuals genotyped per population; **PA/L**: private alleles per locus; **av# alleles**: average number of alleles per locus **Fis**: inbreeding coefficient [42] **Significant Heterozygote deficit (p < 0.05).; **av PIC**: averaged Polymorphism Information Content [74]. **av A_r_**:Averaged allele richness [43]; **He**: expected heterozygosity = average gene diversity [44]; **Ho**: observed heterozygosity; **PL Hd**: percentage of polymorphic loci displaying a deficit in heterozygosity; **D_AS_**= Pairwise distance between alleles. ***I*_A_**: index of association that suggests multilocus linkage disequilibrium if p<0.05.

| Population | #MLG8/N | PA/L ±<br>SE | av# alleles | Fis ±<br>SE | avPIC | av $A_r$ | He ±<br>SE | Ho | %PL<br>Hd | D<sub>AS</sub> ±<br>SE | I<sub>A</sub> | I<sub>A</sub> P-value |
| --- | --- | --- | --- | --- | --- | --- | --- | --- | --- | --- | --- | --- |
| Mataral-D | 18/19 | 0.55 ±<br>0.09 | 5 | 0.052 ±<br>0.03 | 0.57 | 4.51 ±<br>1.91 | 0.63 ±<br>0.07 | 0.6 | 50 | 0.55 ±<br>0.12 | 0.40 | 0.007 |
| Mataral-S | 10/11 | 1.26 ±<br>0.22 | 5.5 | 0.22 ±<br>0.04** | 0.65 | 5.37 ±<br>2.07 | 0.74 ±<br>0.05 | 0.57 | 100 | 0.67 ±<br>0.19 | 1.08 | <0.001 |
| Kirus Mayu | 20/22 | 0.45 ±<br>0.12 | 4.3 | 0.108 ±<br>0.03** | 0.47 | 3.69 ±<br>1.83 | 0.52 ±<br>0.09 | 0.47 | 85.7 | 0.45 ±<br>0.15 | 0.65 | <0.001 |
| Ilicuni | 18/18 | 0.75 ±<br>0.11 | 4.9 | 0.047 ±<br>0.05 | 0.58 | 4.34 ±<br>1.50 | 0.64 ±<br>0.07 | 0.61 | 62.5 | 0.54 ±<br>0.14 | 0.45 | 0.012 |
| 20 de Octubre | 18/22 | 0.42 ±<br>0.13 | 3.6 | -0.047 ±<br>0.07** | 0.37 | 3.06 ±<br>0.96 | 0.44 ±<br>0.06 | 0.46 | 50 | 0.34 ±<br>0.17 | 1.44 | <0.001 |

### Inter-population subdivision and gene flow

Both genic and genotypic Fisher tests for pairwise population differentiation were highly significant for all pairs compared. A hierarchical AMOVA, which evaluated the distribution of genetic diversity, demonstrated that the majority of variation (71.5%) was within individuals, compared to 24.1% and 4.4% among populations and among individuals within populations, respectively. A population neighbor joining (NJ) clustering analysis based on pair-wise genetic distances *D*_AS_ [66] is shown in Figure 1. The NJ topology was consistent with geographical origin and *F*_ST_ distances. The more proximate populations, both Mataral populations, were grouped together and closer to Ilicuni (separated by approximately 60 km), while the more distant populations of 20oct and KM were more closely related.

Estimates of population subdivision based on allele frequency and identity (*F*_ST_) indicate significant genetic differentiation between all populations, with the lowest *F*_ST_ values found between both populations of Mataral and Ilicuni (Table 3). *F*_ST_ values among sylvatic populations spanning a greater geographical distance (e.g. *F*_ST_= 0.27 between Mat-S and 20Oct; 162.5 km) were largely equivalent to those observed between domestic and sylvatic triatomines sampled from a more restricted area (*F*_ST_= 0.22 between Mat-D and KM; 57 km) and between adjacent sylvatic populations (*F*_ST_= 0.22 between Mat-S and KM 102.06 km), suggesting little gene flow occurred among these populations throughout the geographic range considered. The Mantel’s test conducted to evaluate the significance of the correlation between the geographical and genetic distances between population pairs, showed a slight but significant adjustment to an isolation by distance model (Rxy=0.413; P<0.01).

**Figure 1:**
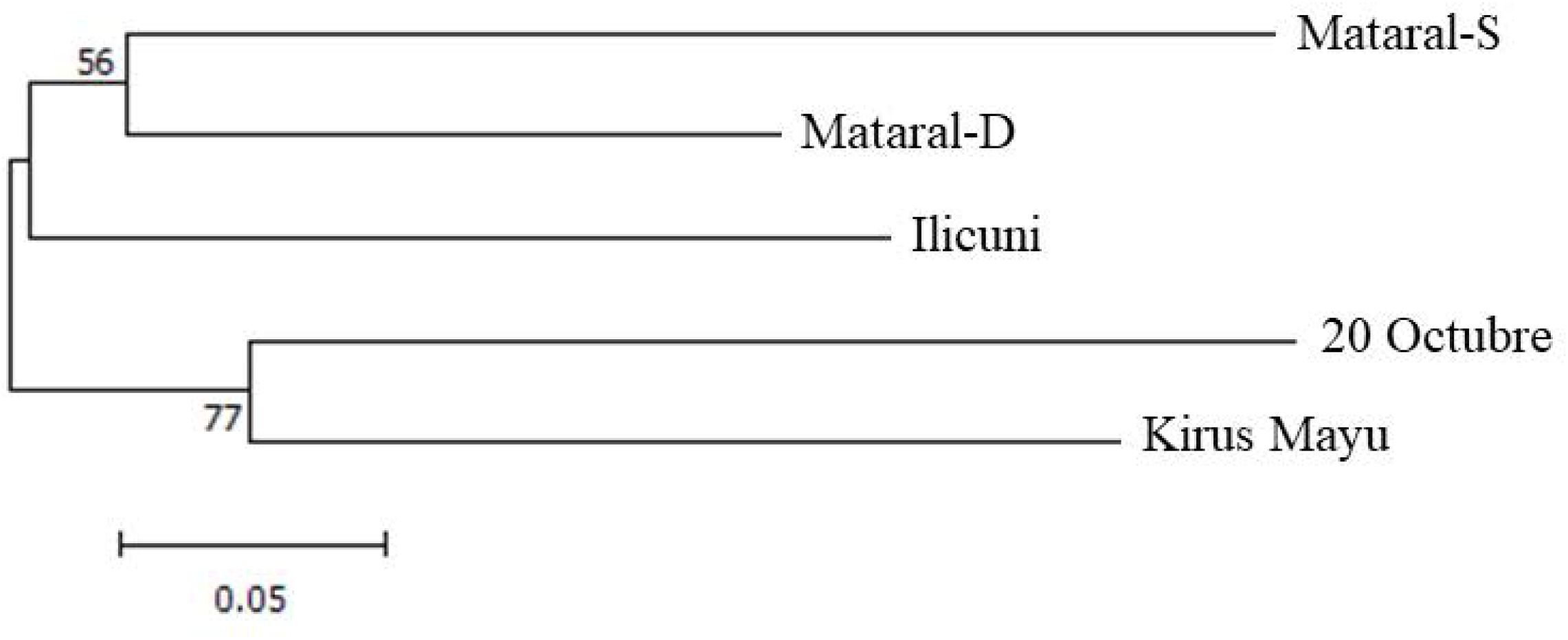
Neighbor joining (NJ) tree based on population pairwise *D*_AS_distances [66] estimated with the MLGs of 8 loci and 1000 replications.

**Table 3:** *F*_ST_ values (geographic distances) in a pair-wise comparisons between populations (** p<0.005)

|  | 20oct | Ili | KM | Mat-S | Mat-D |
| --- | --- | --- | --- | --- | --- |
| 20oct | - | ** | ** | ** | ** |
| Ili | 0.272<br>(151.76 km) | - | ** | ** | ** |
| KM | 0.323<br>(62.92 km) | 0.282<br>(104.63 km) | - | ** | ** |
| Mat-S | 0.272<br>(162.50 km) | 0.144<br>(56.02 km) | 0.223<br>(102.06 km) | - | ** |
| Mat-D | 0.278<br>(159.74 km) | 0.163<br>(56.94 km) | 0.216<br>(99.07 km) | 0.145<br>(3.38 km) | - |

A Bayesian assignment of individual genotypes determined an optimal number of five genetic clusters to describe the dataset (Figures 2 and 3), corresponding broadly to the five populations assigned by capture site. Most individual MLGs clustered significantly with other members from the same geographic location. However, in all sylvatic populations, individuals that could be first generation migrants or of admixed origin were detected (Figure 3).

A hierarchical comparison of alternative clustering models reflects the relative genetic distance among MLGs. This analysis shows the resulting clustering pattern of MLGs from the whole sample when using different K values (i.e. considering alternative numbers of genetic clusters). The results are presented in Figure 3, which shows the closer genetic relationship between both Mataral populations and Ilicuni, as the individuals from those populations cluster together when considering four or three possible genetic clusters to assign the whole dataset.

**Figures 2:**
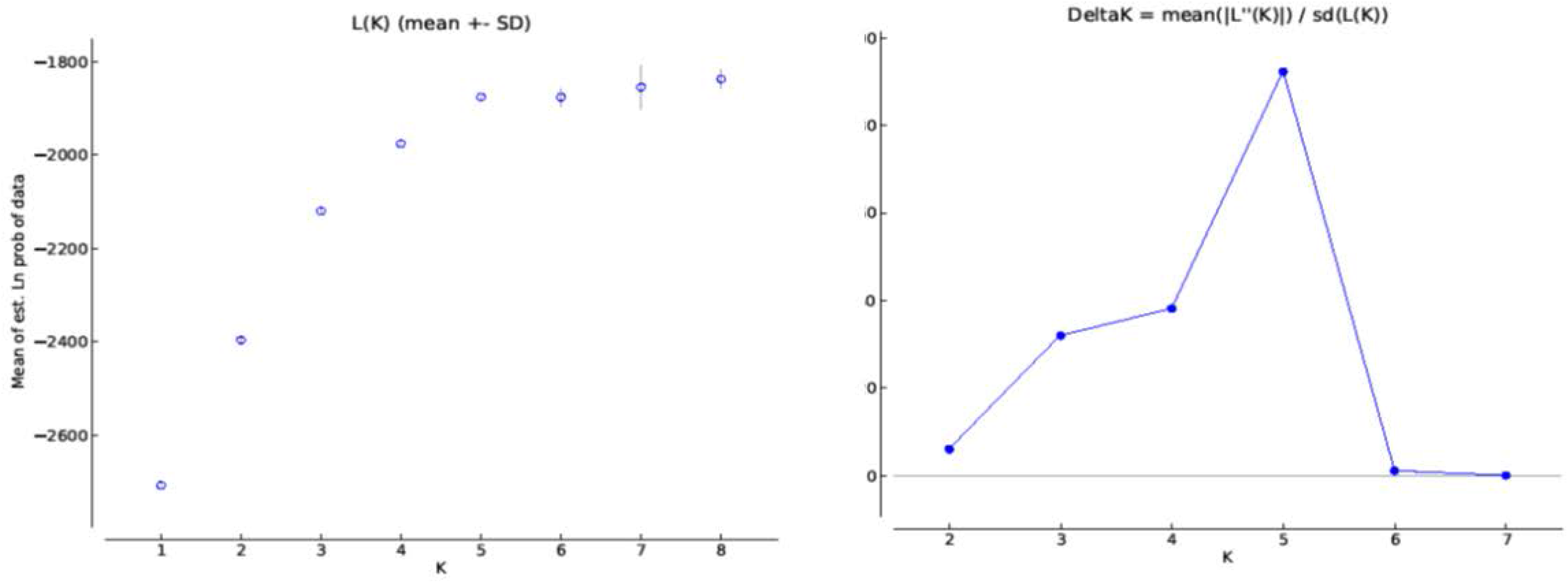
Bayesian cluster analysis results of *T. infestans* populations from sylvatic and domestic areas in Bolivia. 2a) Mean of estimated Ln likelihood of the data (Ln(P)) versus K number of genetic clusters considered [52]. 2b) Graphic representation of Delta K=mean ((|L’(K)|)/SD(L(K)) [55] to evaluate the rate of change in the Ln(P) versus the number of genetic clusters considered (*K*=1-10).

**Figures 3:**
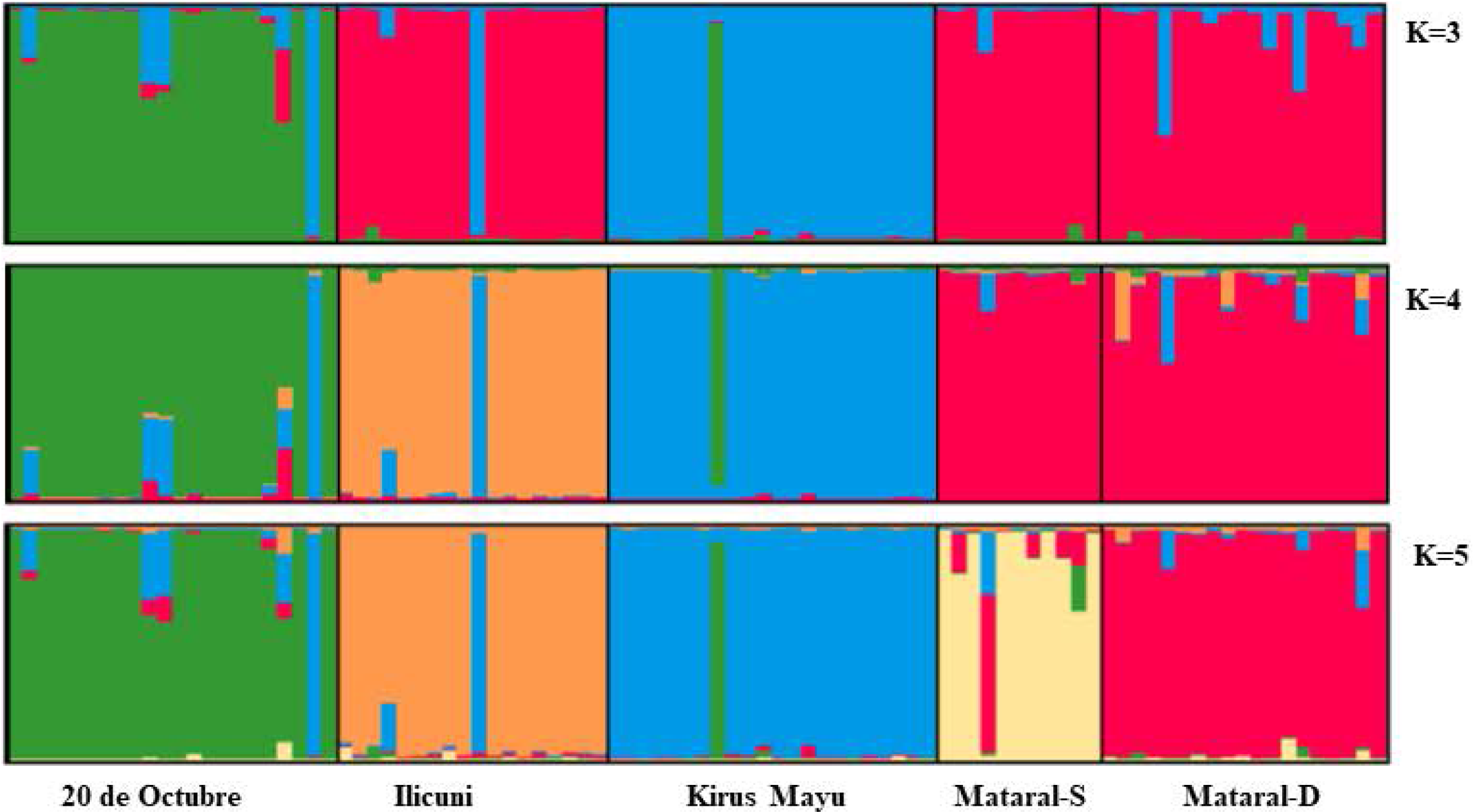
Bar plot representing the individual assignment probability of each MLG to a specific genetic cluster. Each vertical bar represents an individual and the different colors are the probability that each individual was assigned to a particular genetic cluster (K). The hierarchical approach allows for comparison of individual MLG assignment results when considering K=3, K=4 and K=5 genetic clusters.

### Mitochondrial genetic clustering among populations and individuals

A fragment of the cytochrome B gene (cytB) was amplified and sequenced from 74 individuals (20oct=14, Ili=15, KM=15, Mat-S=11 and Mat-D=19), and assembled into a 666 bp alignment. A total of eight haplotypes were observed, including an ambiguous (i.e. heteroplasmid) sequence detected in two individuals from KM (ht). All haplotypes were submitted and are accessible in GenBank (access numbers MH763648 to MH763654). Haplotypes were determined by 15 variable sites that were mostly synonymous except for sites 28 and 625, carried by haplotypes XVI and XLIX (Table 4).

**Table 4:** Variable sites among the cytochrome B (cytB) haplotypes detected in sylvatic populations of Andean *T. infestans* in Bolivia. Highlighted background indicates non-synonymous substitutions. R and Y are the IUPAC nucleotide code indicating ambiguous base position (A/G) and (C/T) respectively. HT: heteroplasmid sequence (i.e. individuals carrying more than one mitochondrial haplotype). Haplotype I was among the first one described for *T. infestans* in Bolivia [75] and found widely distributed in several populations since then. Green indicates the variable sites shared between haplotypes sequenced in this study and those previously deposited in GenBank for Bolivian Andean *T.infestans* populations.

| haplotypes | variable sites (nucleotide positions) |  |  |  |  |  |  |  |  |  |  |  |  |  |  |
| --- | --- | --- | --- | --- | --- | --- | --- | --- | --- | --- | --- | --- | --- | --- | --- |
|  | 28 | 162 | 279 | 402 | 450 | 480 | 526 | 549 | 564 | 576 | 588 | 600 | 625 | 646 | 636 |
| I | G | A | A | A | A | A | C | G | T | A | T | C | G | T | T |
| XXIV | . | . | . | G | G | . | T | . | . | T | . | . | . | . | . |
| XVI | A | G | . | . | . | . | . | . | . | T | . | . | . | . | C |
| HT | A | G | R | . | . | R | . | . | . | Y | . | . | . | Y | C |
| XLIX | A | G | G | . | . | G | . | . | . | C | . | . | A | C | C |
| XXVII | . | G | . | . | . | . | . | . | . | T | . | . | . | . | C |
| XLV | . | G | . | . | . | . | . | . | . | T | . | . | . | . | C |
| XLVI | . | . | . | . | . | . | . | A | . | T | C | . | . | . | . |
| XLVII | . | . | . | . | . | . | . | A | C | T | C | . | . | Y | C |

The most common mitochondrial genotype was XVI, identified in 21 isolates from 20oct, Ili and KM, but absent from both Mataral populations (Table 5, Figure 4). Mat-D and Ili displayed the highest haplotype diversity (Hd=0.65 and 0.59, respectively), and were the populations that also presented the most diverse haplotypes (π=0.0056 and 0.0044, respectively) (Table 5). Both Mataral populations presented private unique haplotypes (XLV in Mat-D and XLVI and XLVII in Mat-S, respectively). Only one of the five haplotypes observed in Mat-S and Mat-D was shared between the two populations (XXIV).

**Table 5:** Cytochrome B (cytB) haplotype distribution per population and diversity parameters. **Hd**= Haplotype diversity, **π** = nucleotide diversity, SD = standard deviation. **ht** = heteroplasmid

| Population | XXVII | XXIV | XVI | XLV | XLVI | XLVII | XLIX | ht | # cytB<br>seqs | Hd ± SD | $\pi$ ± SD |
| --- | --- | --- | --- | --- | --- | --- | --- | --- | --- | --- | --- |
| Mat-D | 7 | 9 |  | 3 |  |  |  |  | 19 | 0.649 ± 0.061 | 0.00437 ± 0.00033 |
| Mat-S |  | 1 |  |  | 9 | 1 |  |  | 11 | 0.455 ± 0.17 | 0.00241 ± 0.0011 |
| KM | 1 |  | 2 |  |  |  | 10 | 2 | 15 | 0.41 ± 0.154 | 0.00312 ± 0.0011 |
| Ili |  | 6 | 8 |  |  |  | 1 |  | 15 | 0.59 ± 0.077 | 0.00563 ± 0.00097 |
| 20oct |  |  | 11 |  |  |  | 3 |  | 14 | 0.363 ± 0.13 | 0.00272 ± 0.00098 |

**Figure 4:**
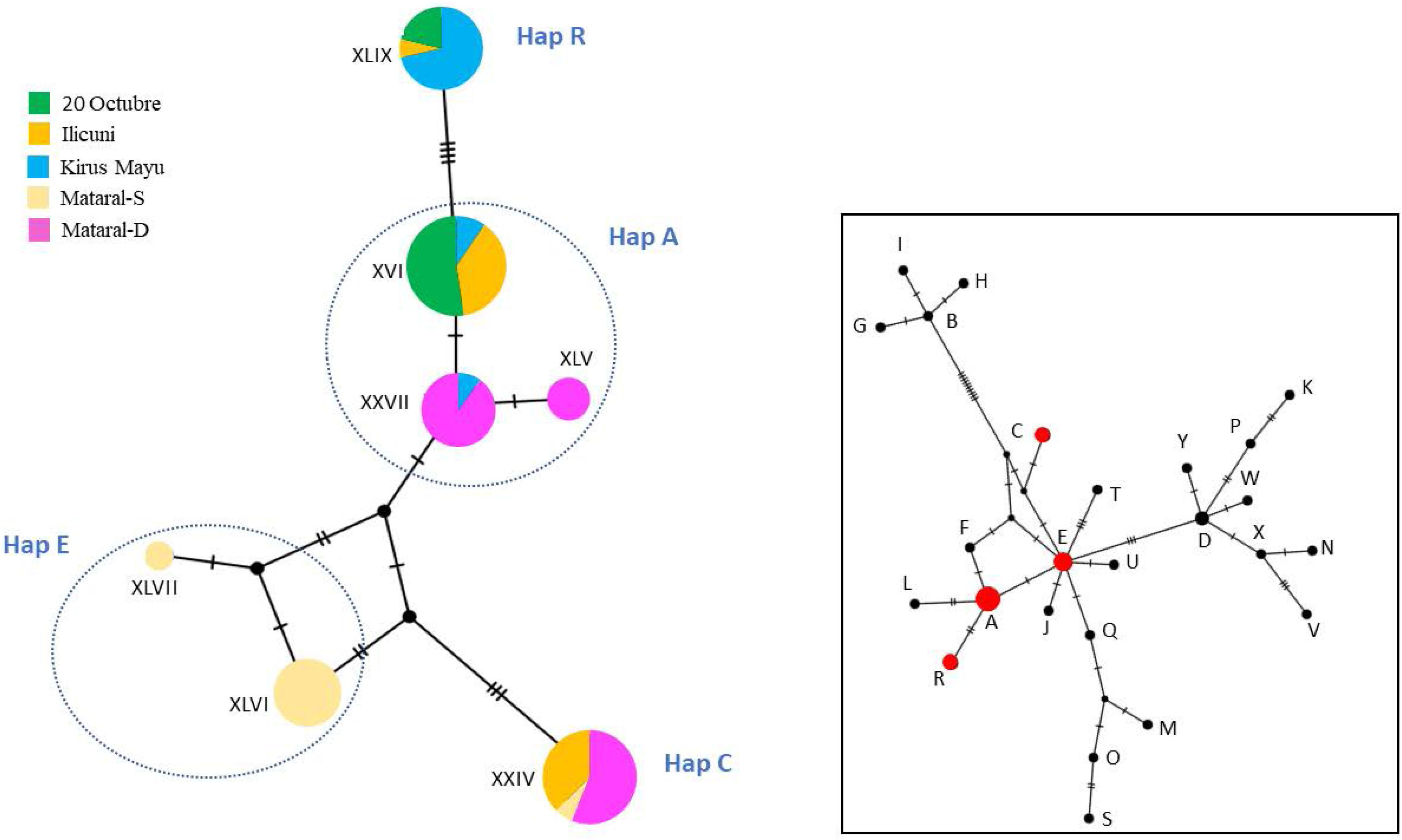
Left: haplotype network based on a median-joining model, constructed from sequences identified in this study. The size of the nodes is proportional to the abundance of each haplotype. The colors indicate the population of origin. The distance among haplotypes is represented by the number of mutational steps between them. Right: haplotype network including 25 additional Bolivian Andean *T. infestans* haplotypes (labelled A-Y) [15, 62]. Trimmed sequences from this study corresponded to four haplotypes previously reported: A, C, E and R (shown in red).

The distribution of mitochondrial haplotypes across populations corresponded broadly to the geographical distances between populations, with KM sharing two genotypes with 20oct (XVI and XLIX); likewise Ili and both Mataral populations contained more closely-related cytB sequences. The phylogenetic tree and network analysis, clustered haplotypes into two groups based on the substitution A/G in nucleotide position 162 (Table 4). One cluster contained haplotypes XLVI and XLVII (only detected in Mat-S population) with XXIV, found in both Mataral populations and Ilicuni. The remaining haplotypes were found in all but the Mat-S population (Figure 4).

In order to interpret the results of this study in the context of known phylogeographic patterns of *T. infestans* populations in Bolivia, we built a haplotype network with previously reported haplotypes for Bolivian *T.infestans* [15, 62]. As the sequences previously reported were only 388bp long, the haplotypes we characterized were trimmed to that length for comparison. Consequently, some of our trimmed haplotypes were indistinguishable from each other within the fragment length considered, and were pooled into four haplotypes previously reported in Andean Bolivian populations: A, C, E and R [15] which are shown in red in Figure 4.

Remarkably, the haplotype network built with sequences from previous studies of Bolivian *T. infestans* (Fig. 4) showed that the haplotypes found in Mataral-S correspond to a central node (i.e. ancient), haplotype E, composed of populations from the “Central Andean area” collected in two locations in Cochabamba, Mataral [15] and Chuquisaca [67]. Another haplotype found in Mataral-S which has also been found in Mataral-D and Ilicuni (haplotype XXIV from this work), corresponded to the previously reported haplotype C, a rare divergent haplotype from most of the others reported and considered a derivate from haplotype E [15].

In this comparison, the three haplotypes found in 20 Octubre, Kirus Mayu, Ilicuni and Mataral-D populations correspond to that described as the most abundant haplotype A, previously detected in specimens from “Northern and Central Andean areas” [15]. Finally the haplotype XLIX found in 20 Octubre, Kirus Mayu and Ilicuni correspond to haplotype R, considered a derivate of haplotype A [15]. Haplotypes A, R and C had also been previously reported in domestic populations [67].

### *Kdr* screening

Among the sixty-seven samples screened for two mutations in the *kdr* gene, related to insecticide resistance in *T. infestans*, only one individual from Mataral-S presented the resistant allele variant for the site L1014. No individual carrying the mutation L9251 was detected.

## Discussion

Through high resolution genotyping of five Andean *T. infestans* populations, we provide insights into natural population genetic structures and the relationships between triatomine genetic diversity and insecticide resistance. All populations were genetically distinct, which correlates with previously described heterogeneity in insecticide susceptibility profiles [24]. The pattern of interpopulation differentiation was consistent with a significant isolation-by-distance (IBD) model, with geographically closer populations being more genetically-related. This observation was supported by both mitochondrial and nuclear microsatellite markers. The genetic pattern of allelic similarity reflected possible gene flow and/or secondary contact between domestic and sylvatic Mataral populations. This observation furthers supports previous reports of dynamic geneflow between sylvatic and domestic populations in the Bolivian Andes [15, 36, 62].

Both populations from Mataral were characterized by considerably higher genetic diversity compared to the other sylvatic populations evaluated in this work, and between the Mataral populations, genetic diversity and substructure were higher for the sylvatic population. Furthermore, high levels of homozygosity and numbers of private alleles per locus suggest that Mataral-S may be ancestral to the other sylvatic populations evaluated here. This observation is independently supported by the occurrence of two mitochondrial haplotypes within this population that correspond to haplotype E, which has been previously reported as an ancient haplotype among other Andean *T. infestans* populations [15].

The populations 20oct, KM and Mat-S presented significant deviations from HW equilibrium, and all due to significant heterozygote deficits. As the samples were arbitrarily grouped per community, the significant Fis values are likely attributable to the population being comprised of admixed individuals from multiple, independent non-panmictic founding populations (Wahlund effect). This would also account for the higher linkage disequilibrium detected in these populations. High standard deviations associated with *D*_AS_values further supports the existence of intra-population sub-structuring. Alternatively, this genetic pattern could evidence that some members have undergone long-term inbreeding, following historical isolation.

The bioassays performed in these populations demonstrated three different levels of altered susceptibility [24]. High levels of decreased susceptibility in Mataral-D and Mataral-S, moderate levels of decreased susceptibility in Kirus-Mayu and 20 de Octubre and limited altered susceptibility in Ilicuni (Table 1). The RRs obtained for these populations are not as high as values from other areas that have presented difficulties for vector control (i.e. RR>2000 El Juramento, Guemes, Chaco, Argentina [26] or RR_50_=541.6, Tierras Nuevas, Tarija, Bolivia [68]). However, in all but Ilicuni, the RR values are above the level classified as resistant by PAHO guidelines RR>5 (fivefold higher than reference populations) [30, 69]. Those guidelines also attribute altered susceptibility in field populations presenting RR<5 to individual variability, which is not enough to cause resistance at the population level. As previously discussed [25, 70], elevated RR in a sylvatic population might result from at least three non-mutually exclusive hypotheses:

a. existence of naturally altered (lower) susceptibility to insecticides
b. development of resistance resulting from exposure to insecticides used in agriculture and vector control campaigns
c. contact and exchange of genetic variants between sylvatic and domestic resistant populations

In the context of this study, the decreased susceptibility to deltamethrin observed in all populations may represent a shared ancestral trait, which has undergone further intensification in Mataral following insecticidal treatment, particularly for the domestic population, with possible geneflow between the two Mataral populations, including the dispersal of bugs with altered susceptibility to Pyrethroids. In this scenario, the intermediate RRs of 20 de Octubre and Kirus-Mayu would have been maintained by moderate selection pressure, likely the indirect effect of agricultural spraying; the difference between them could be explained by intrinsic intra-species variability, consistent with the distinct genetic patterns observed. However, the minimally altered susceptibility observed in Ilicuni would imply that this trait had been almost lost in this population in the absence of selection. Alternatively, insecticide resistance could have been acquired independently in all populations following different histories of insecticide exposure, with population RR directly corresponding to the intensity of selection each population has undergone. Further characterization of the ecological context of the population in this area is necessary to test these hypothesis or propose alternative explanations to observed the genetic pattern.

Insecticide-based interventions are expected to impact vector population structure and levels of genetic diversity. Selective bottlenecks imposed by insecticidal pressure are likely to result in both reductions in genetic variability and population size, such that a small proportion of survivors, possessing the requisite advantageous resistance genes, survive and reproduce (i.e. diversification that would be reflected in a genetic pattern as multiple independent “island effects”). In general, insecticide resistant populations are expected to be less genetically diverse than a wild-type population, as they have likely undergone a period of selective pressure (i.e. insecticide exposure). And within this context, domestic populations are assumed to have been the subject of greater selection pressures, compared to their sylvatic counterparts and therefore be less diverse.

However, our study results challenge these traditional paradigms and may instead be explained by an alternative hypothesis for genetic diversification. In a given location, a population may undergo a selection process from insecticide exposure independent of neighboring triatomine populations [71]. A previous work comparing the genetic profiles of populations under different vector control pressure [33] described populations within a community recovering from a recent massive spraying as much more genetically structured and diverse than populations that were not under systematic vector control pressure. The authors proposed that the distinct observed genetic pattern reflected the cumulative effect of migration among adjacent populations for several generations, which generated a pattern of genetic “homogenization” even among populations located geographically far apart. In contrast, populations recovering from a pool of survivors of insecticide spraying were observed as discrete genetic entities at the individual capture site (i.e. one household). Pooling together such individuals from these “genetically discrete” populations with bugs from other sites, for a random sampling of a given area, (i.e. at the community level, pooling individuals from different houses in the same sample set), would result in a highly genetically diverse metapopulation, that is actually composed of several subpopulations, with genetic substructure and gene flow restrictions within. In this study, the elevated genetic diversity of both Mataral populations could represent the assemblage of individuals from distinct genetic units, as they have been under higher selective pressure than the sylvatic populations from 20 de Octubre, Kirus-Mayu and Ilicuni, located further away from the domestic area subjected to vector control. Great genetic diversity has also been reported in several domestic non-Andean *T. infestans* populations from the Gran-Chaco area [35, 72], of which many had been under extensive vector control pressure.

More importantly, the existence of genetically-diverse populations displaying high levels of altered insecticide resistance has potential implications for longer-term efficacy of control interventions, such that the likelihood of individuals surviving future selection events is greater when the original population is composed of more unique genetic variants. It is noteworthy that three haplotypes found in the sylvatic populations analyzed here (A, C and E) were also identified in domestic populations in the department of Cochabamba [67], indicative of household colonization by sylvatic populations in the Bolivian Andes, a pervasive and contemporary process.

Regarding the resistance mechanism(s) underlying the altered susceptibility patterns in Mataral, analyses of P450 monoxygenases and 7-CP esterases [24] showed a lack of association between enzymatic activities and susceptibility ratios, indicating that these enzymes were not responsible for modified susceptibility in these sylvatic populations. In this work, we evaluated the presence of two *kdr* mutations and found one individual from sylvatic Mataral carrying the mutation L1014 [63]. This result indicates that although not likely responsible for the mechanism altering insecticide susceptibility in these populations, the presence of the mutation in this area, represents a high risk for selection under pressure and potential development of higher resistance intensities [73]. Future work to characterize the underlying metabolic mechanisms of resistance in these populations is warranted.

## Conclusions

Sylvatic populations of *Triatoma infestans* represent a challenge for vector control. These populations are not targeted by control activities and could play a key role in post-spraying house re-infestation. The results presented here suggest that there might be an ancient trait(s) that confers resistance in *T. infestans* sylvatic populations that are capable of invading domiciles. These observations emphasize the need for stronger entomological surveillance in the region, including early detection of house invasion, particularly post-spraying, monitoring for resistance to pyrethroids and the design of integrative control actions that consider both sylvatic foci around domestic settings as well as the bug dispersion dynamics.

## Declarations

### Ethics approval and consent to participate

Not applicable.

### Consent for publication

Not applicable.

### Availability of data and material

DNA sequences obtained in this study are available in Genebank.

### Authors’ contributions

PLM, PSH, CVV conceived the idea and designed the work. PLM carried out the laboratory work. PLM and LAM performed the analysis and took the lead in writing the manuscript. All authors provided critical feedback for the manuscript and contributed to the conclusions of the work. All authors read and approved the final version of the manuscript.

## Acknowledgements

No applicable

